## supplementary_figures for "Cerebrospinal fluid proteome maps detect pathogen-specific host response patterns in meningitis"

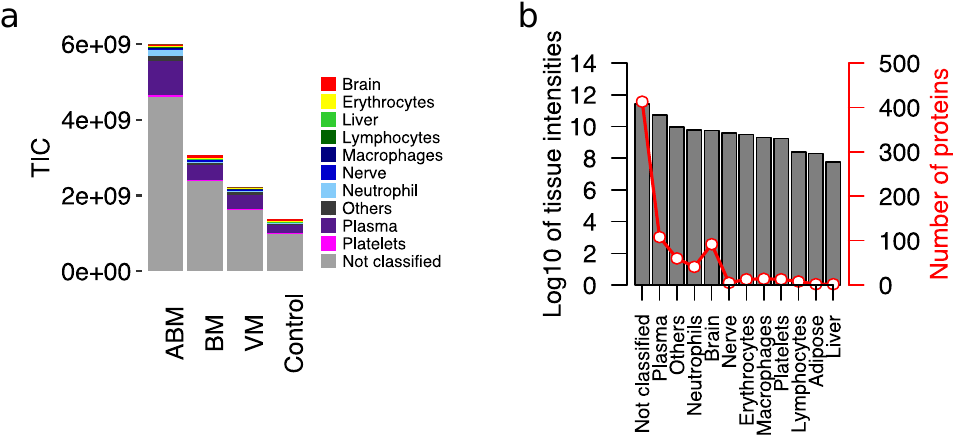

**Fig. S1.** Overview of average protein content and intensities. (Related to figure 2.) (a) Average total ion currents (TIC) for ABM, BM, VM and control groups are presented, and the contribution of each tissue group is shown with color. (b) A dual-axis graph showing the average intensities for each 12 tissues (left y-axis) and the number of proteins associated to each tissue (right y-axis).

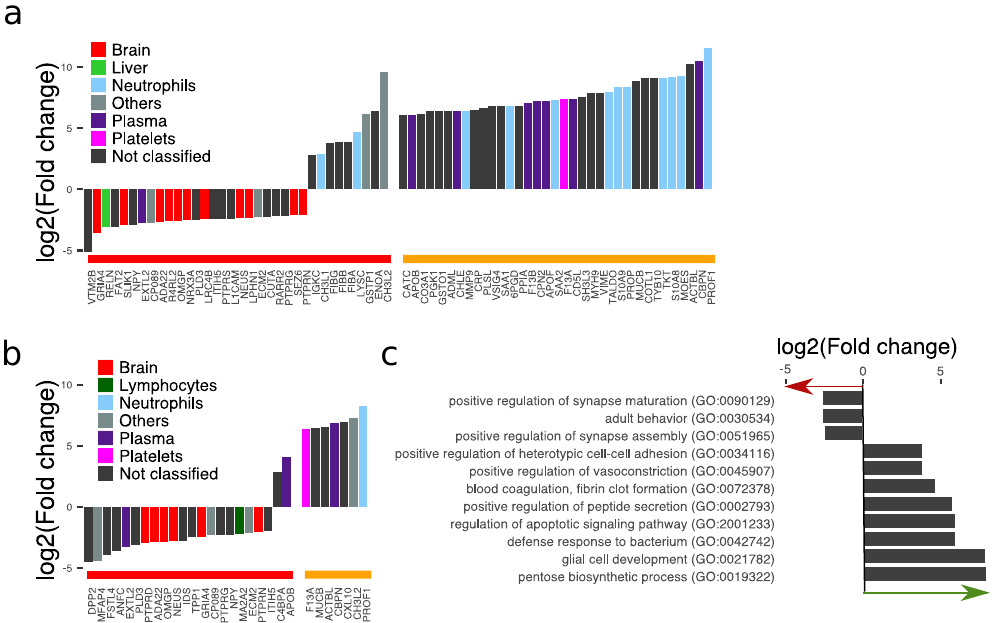

**Fig. S2.** Analysis of differences in the CSF of patients with meningitis. (Related to figure 2.) (a-b) The tissue-assignment, protein abundances and identities of proteins with statistical significance (Benjamini-Hochberg-corrected p-value ≤ 0.05 and log2 fold change ≤ -2 and ≥ 2) are shown and labeled with red horizontal line. Statistically non-significant proteins with high fold change of ≥ 64 were considered of biological interest and labeled with yellow horizontal line. The data is shown for ABM in (a) and for VM in (b). (g) Proteins selected for ABM were annotated by gene ontology (GO) terms. The average fold change for all proteins included in each term are plotted to show the direction of change compared to controls.

File: Supplementary_table1.xlsx

Table S1. Cross-referencing the tissue assignments based on Malmström, et al. (unpublished work). The tissue assignments based on Malmström, et al. (unpublished work) for proteins in figures 3F-I, 4C, 5C and 5F and supplementary figures 2A-B matched and compared to publicly available and published protein tissue assignment repositories.

File: Supplementary_table2.xlsx

Table S2. The full data generated from 112 data-independent acquisition MS-runs (ABM: n=25, BM: n=7, VM: n=21 of which TBE: n=5 and controls: n=49). The data was manually curated to remove immunoglobulin variable chain proteins.

File: Supplementary_table3.xlsx

Table S3. The full data generated from the data-independent acquisition MS-runs from the longitudinal study. The cohort consists of longitudinal samples collected from 6 ABM patients (6 original samples used in this study and additional 14 longitudinal samples) and from 4 patients with subarachnoidal hemorrhage (SAH, 22 longitudinal samples). The data was manually curated to remove immunoglobulin variable chain proteins.
